# Design to Data for mutants of β-glucosidase B from *Paenibacillus polymyxa*: Q22T, W123R, F155G, Y169M, W438D, V401A

**DOI:** 10.1101/2019.12.23.887380

**Authors:** Chengrui Hou, Paul Smith, Joyce Huang, Jason Fell, Peishan Huang, Ashley Vater, Justin B. Siegel

## Abstract

A key goal of protein engineering is to accurately model the stability and catalytic activity of enzymes. However, the limitations of functional predictive abilities pose a major challenge for modeling algorithm design, and can be attributed to the lack of large data sets quantifying the functional properties of enzymes. Here, the thermal stability (T_M_) and Michaelis-Menten constants (*k*_cat_, K_M_, and *k*_cat_/K_M_) of six new variants of the β-glucosidase B (BglB) protein are quantitatively characterized. Molecular stability of the enzyme variants were hypothesized using the Foldit software and BglB was synthesized in *E. coli* cells. Testing was done through a colorimetric kinetic assay and thermal stability fluorescence-based protein unfolding assay. Results from the assays suggest that all mutations, with the exception of variant Y169M, all experienced reduced catalytic efficiency compared to the wildtype. Assay results indicate that variant W123R is more thermally stable compared to the wildtype, while the differences in thermal stability between the other variants, and the wildtype are negligible. The collected thermal stability and catalytic efficiency data has been added to a data set with the aim of improving Rosetta algorithms for modeling and predicting the functional interactions between biomolecules through a machine learning algorithm and facilitate the precise engineering of protein catalysts.

## Introduction

Enzymes are macromolecular biocatalysts that efficiently facilitate biological reactions in cells. A goal of enzyme engineering is to develop stable, catalytically proficient enzymes for reactions that are not catalyzed by naturally occurring biocatalysts.^1^ For example, bio-engineered catalysts are used in the generation of livestock feeds from the decomposition of cellulosic biomass.^2^ For most applications of biocatalysts, the temperature of denaturation of an enzyme and its quantitative ability to catalyze reactions is key to achieving the desired effects.^3^ Although it has been generally thought that there are tradeoffs between the thermal stability and catalytic efficiency of enzymes^4^, previous studies have found that for β-glucosidase B (BglB), a well-studied enzyme from *Paenibacillus polymyxa* with considerable functional ubiquity, thermal stability and kinetic efficiency can be independently designed.^5^ Currently, the enzyme design community has been challenged by the lack of large, quantitative standardized data sets of functional variant properties. A previous study of the 100 BglB variants revealed that current algorithms do not enable robust and accurate predictions of kinetic parameters. However, machine learning analysis indicated that algorithms, which predict function based on calculated structural features, could be developed with more training data.^6^

Here, six single amino acid variants of BglB are quantitatively characterized and compared with computational predictions through a design-build-test methodology. The mutants are first designed and visualized using Foldit,^7^ and predictions for enzyme stability are quantified through a measure of total system energy score (TSE).^8^ As modeled, lower TSE indicates likeliness for increased stability of the enzyme in complex with the substrate.

Based on the TSE predictions associated with mutations in this study, we hypothesized that the thermal stability of the variants would be overall reduced as compared to the wild type. The physio-chemical molecular interactions - such as hydrogen bond formation - that we observed in Foldit suggest non-generalizable, variant-specific effects on the catalytic efficiencies of the variants.

## Methods

### Mutant Design

Foldit Standalone^7^ was used to design and model six variants of BglB as previously described.^6^ The six mutational changes were scored by the Rosetta energy function and a total system energy score (TSE) is given. All variants were chosen with no greater than five TSE change between the wild type and variant scores to ensure proper folding of the enzyme.^9^

**Figure 1.**
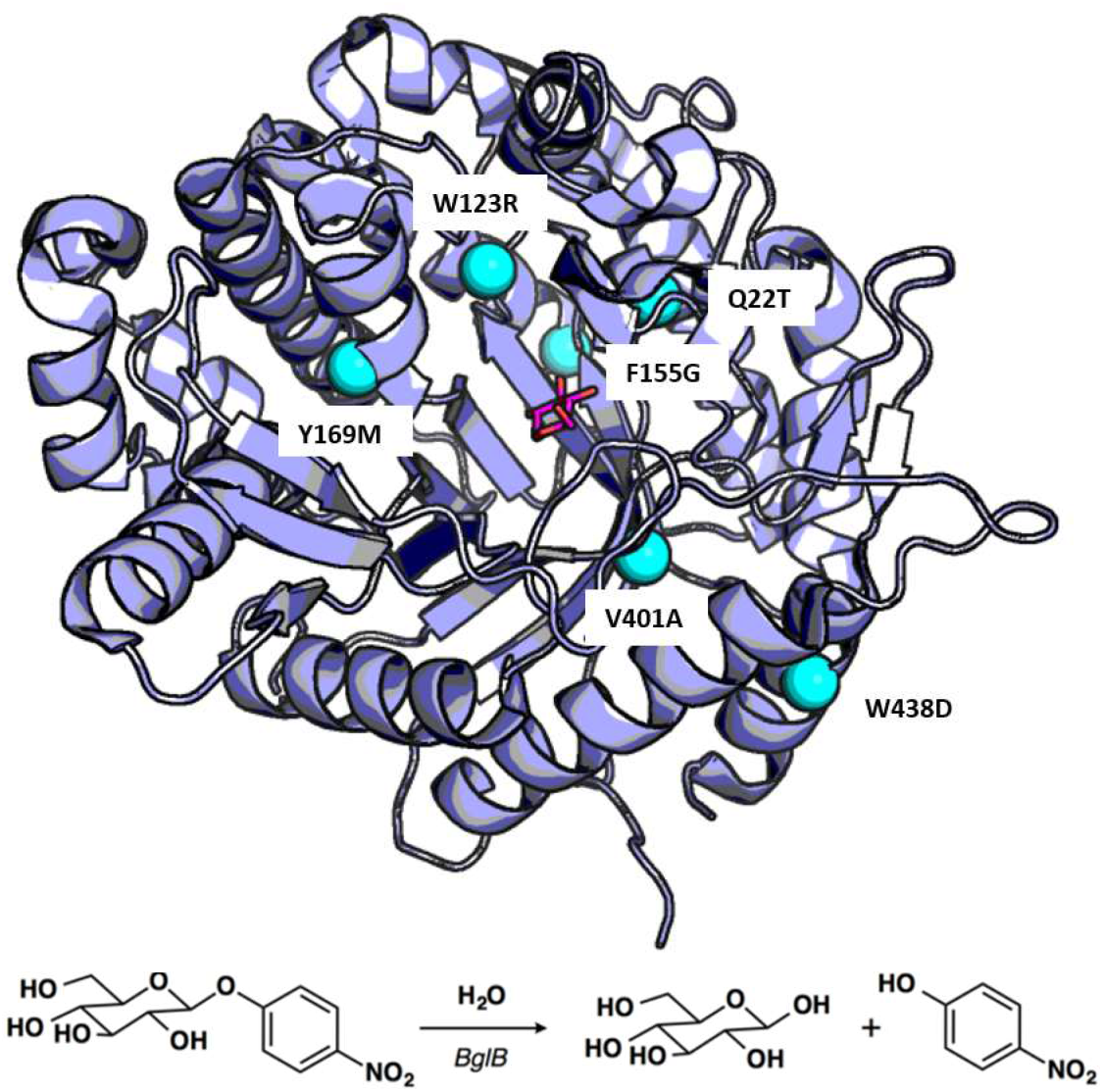
Overview of the BglB and reaction used to determine functional properties of individual variants. PyMOL rendering^10^ of modeled BglB in complex with 2-deoxy-2-fluoro-a-D-glucopyranose (pNPG) showing the six positions selected for mutation in this study (cyan spheres). Below, a reaction scheme of the hydrolysis of pNPG by BglB used to determine functional T_M_ and kinetic parameters *k*_cat_, K_M_, and *k*_cat_/K_M_.

### Molecular cloning and mutagenesis

Standard Kunkel Mutagenesis procedure^11^ was used to generate mutant plasmids used in downstream workflow experiments. Individual plasmid constructs were verified by Sanger sequencing (Eurofins Genomics).

### Protein Production

The sequence verified BglB mutant DNA were transformed, grown, and expressed using the previously described method.^5^ Briefly, following induction using isopropyl β-d-1-thiogalactopyranoside (IPTG), the cells were lysed and the proteins were purified using immobilized metal ion affinity chromatography. Total protein yield was determined by A280 using a BioTek® Epoch spectrophotometer. Protein purity was assessed by SDS-PAGE.

### Determination of Michaelis-Menten kinetics and thermal stability

The kinetic characterization was done in the same manner as the previous study.^6^ Production rate of 4-nitrophenol from the substrate p-nitrophenyl-beta-D-glucoside (pNPG) was determined and recorded through A420 spectrophotometry assay for 1 hour. *K*_cat_ and K_M_ are determined by fitting the data to the Michaelis-Menten kinetics equation.^12^

The thermal stability fluorescence-based protein unfolding assay quantifies the unfolding parameter T_M_, the temperature at which 50% of the protein unfolds. The thermal stability (T_M_) for the variants were determined using the Protein Thermal shift (PTS)™ kit made by Applied BioSystem ® from Thermo Fisher. Following the standard protocol by the manufacturer, purified proteins were diluted from 0.1 to 0.5 mg/mL and fluorescence reading was monitored using QuantaStudio™ 3 System from 20 °C to 90 °C. The T_M_ values were then determined using the two-state Boltzmann model from the Protein Thermal Shift™ Software 1.3 by Applied BioSystem ® from Thermo Fisher. Differences between variant T_M_ parameters and the wildtype are tested using two-sample t-tests. Pearson correlation (PMCC) was used to correlate the predicted change in TSE to measured T_M_ values of variants.

## Results

### Protein expression and purification

Of the six mutant proteins produced, all six expressed and purified as a soluble protein. These variants appeared visibly through SDS-PAGE analysis as shown in Figure 2.

**Figure 2.**
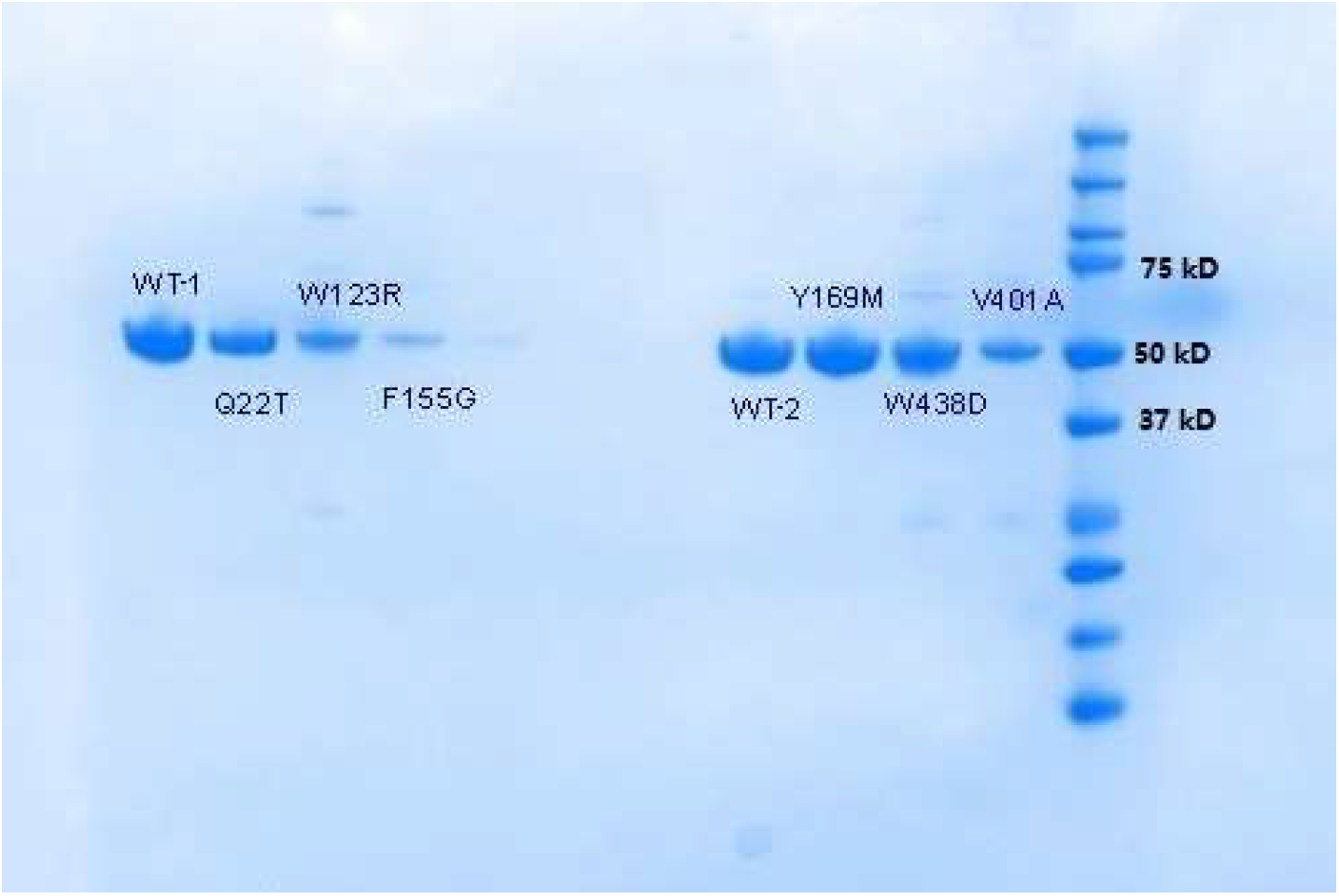
SDS-PAGE gel analysis of wild type and variant enzymes. The purity of protein samples were evaluated using SDS-PAGE. Sample bands formed at 50 kD, indicating BglB enzyme purity. Variants shown from left to right: Q22T, W123R, F155G, Y169M, W438D, V401A

### Mutant Groupings

The six variants of BglB are categorized into three groups based on the intended purpose of our mutations. The groupings are as follows: Group 1 consists of variants that play a direct role in interacting with the substrate: W123R and Q22T. Group 2 consists of variants that indirectly affect the overall hydrogen bonding network in the active site, F155G, V401A, and Y169M. Group 3 consists of variants on the periphery of the enzyme, W438D.

**Figure 3.**
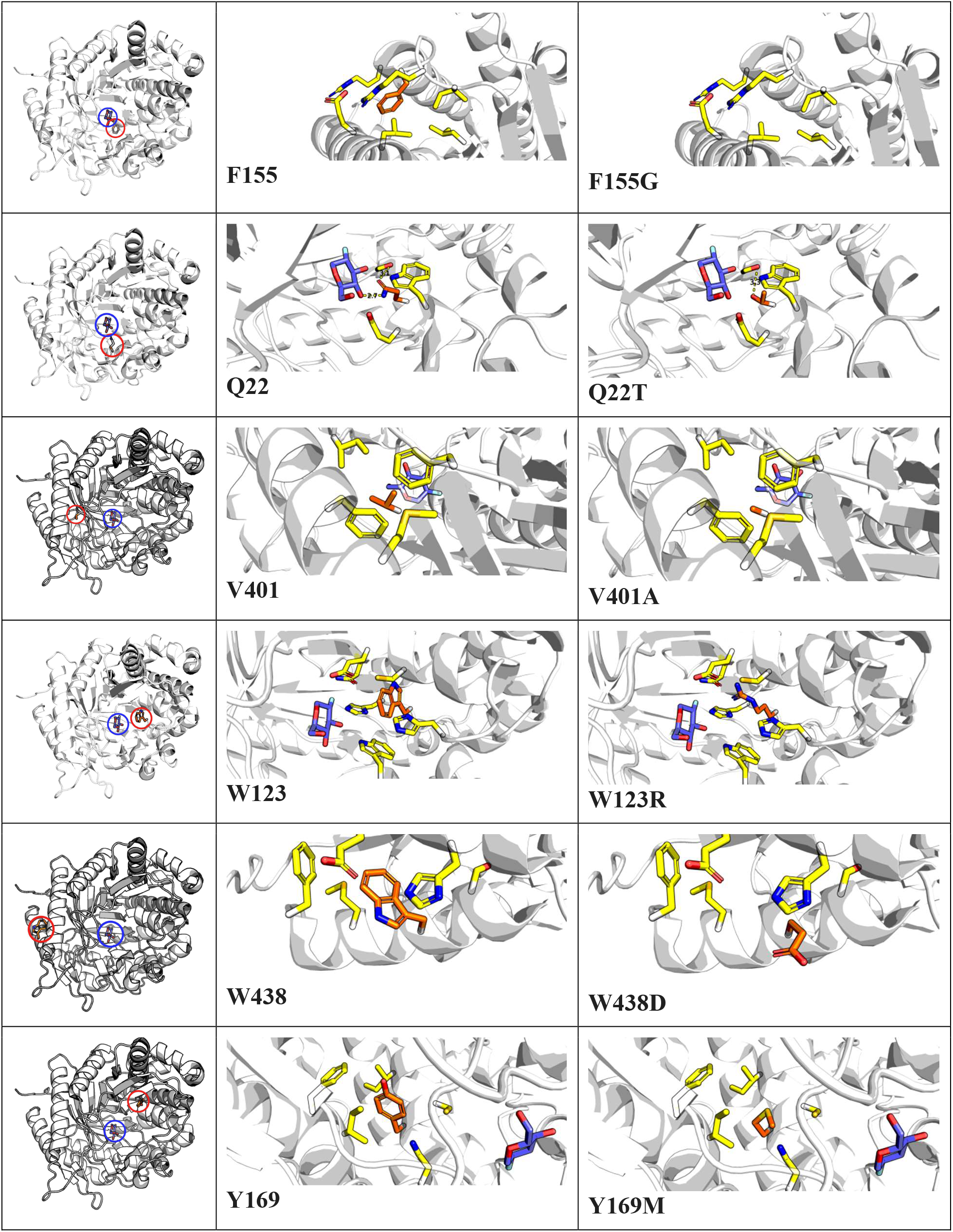
Structural Representation of Variants. The relative locations of mutation sites and molecular interactions of each mutation described in this study are shown. Each mutation is illustrated with three PyMOL^10^ generated images showing site location (mutation site and ligand highlighted in red and blue, respectively), molecular interactions of native amino acid, and molecular interactions of variant amino acid from left to right respectively.

### Kinetic Assay

For the wild type BglB, the *k*_cat_/K_M_ value was determined to be 6.89 and 7.89 min^−1^M^−1^ for the biological replicates, which fall within a two-fold range of the previously established mean *k*_cat_/K_M_ for BglB.^6^ The Q22T and W123R mutations exhibited little catalytic activity, with *k*_cat_/K_M_ values less than 0.04% of the wildtype. The BglB enzyme with F155G, V401A, Y116M and W438D mutations exhibited modestly reduced activity as compared to the wildtype, ranging from 44.16% (F155G) to 14.9% (W438D).

**Figure 4.**
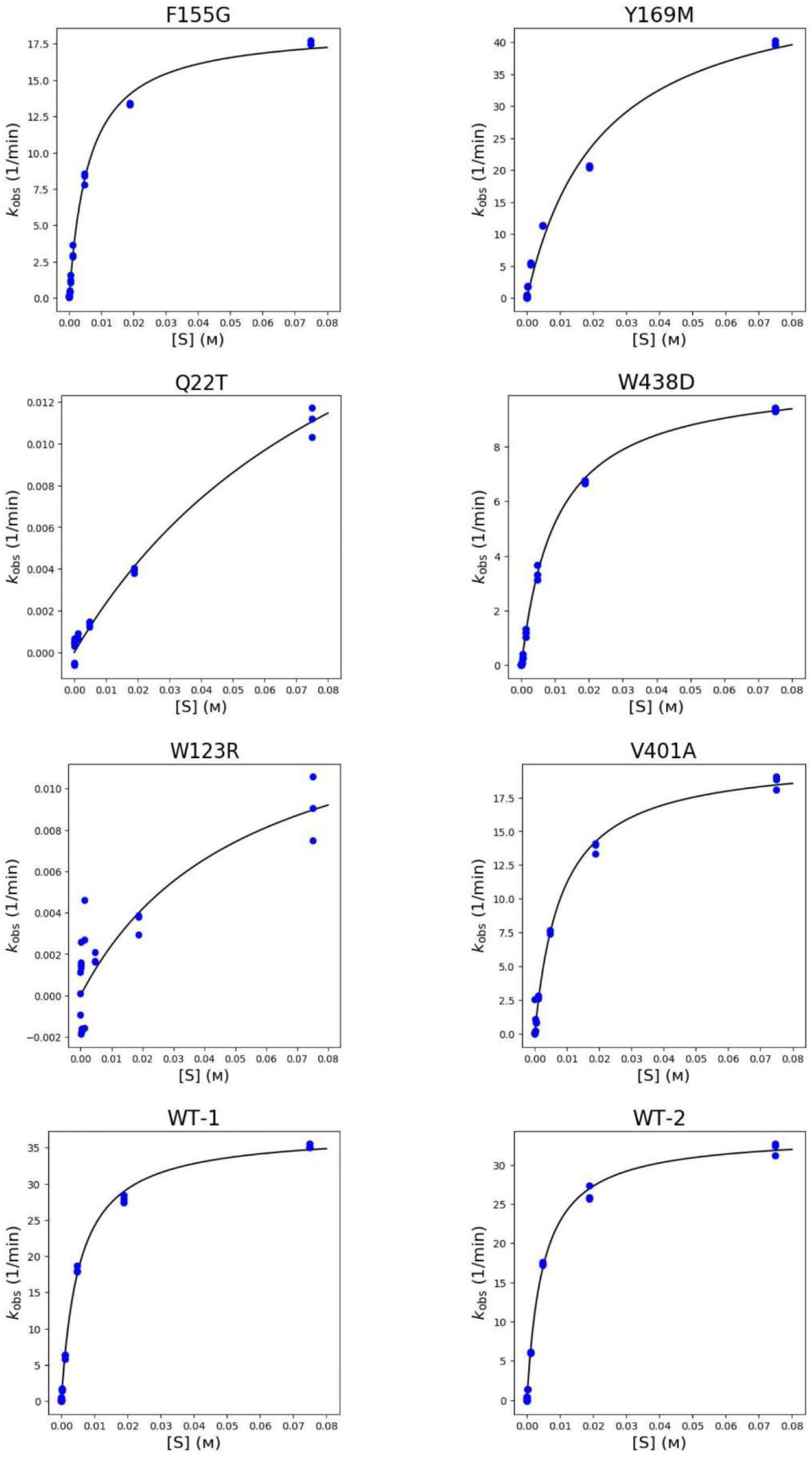
Kinetic Assay Graphs. Variants F155G, W123R, Q22T were tested with WT-1; Variants V401A, Y169M, W438D were tested with WT-2. Two panels of variants are shown, sorted in top-down order by decreasing *k*_cat_. Concentrations of pNPG are plotted on the horizontal scale and *k*_obs_ values are on the vertical scale, with units of M, min^-1^, respectively.

### Thermal Stability Fluorescence-based protein unfolding assay

The mean distribution of T_M_, the temperature at which half the protein molecules denatures, of wildtype BglB is determined to be 44.54 °C, which is in agreement with previously observed T_M_.^9^ The T_M_ of the Q22T mutation did not differ significantly from that of the wildtype BglB (two sample T test, p = 0.70). There was a significant difference between the mean T_M_ of W123R and the wildtype. (p = 0.07).

**Figure 5.**
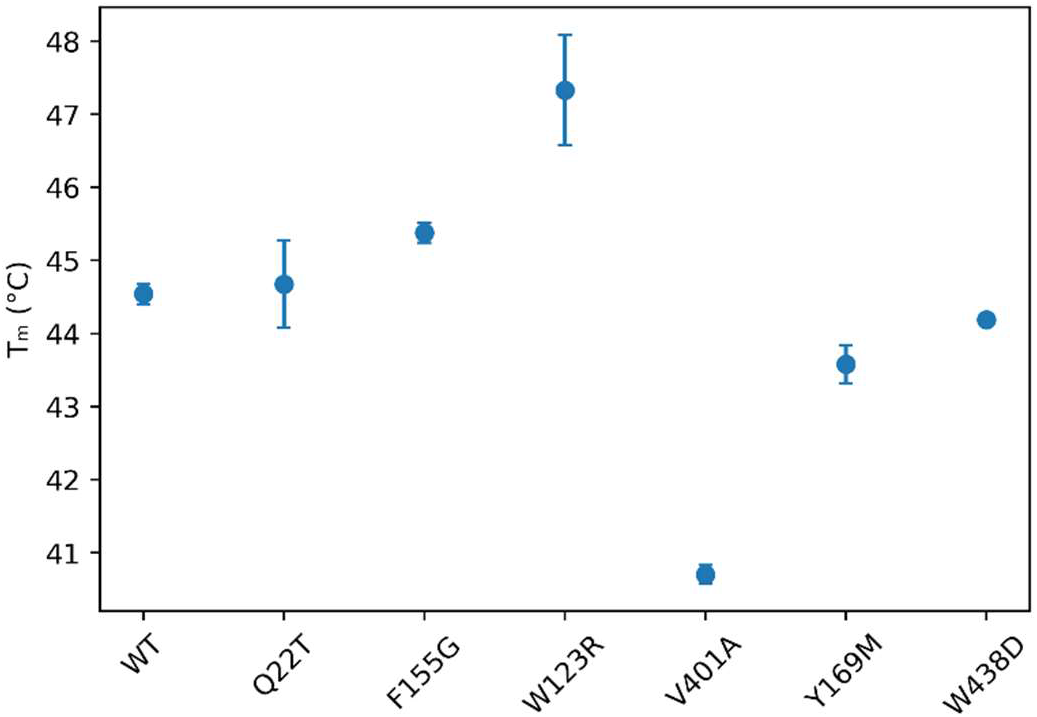
Distribution of Thermal Assay Data. T_M_ values are represented in units of degrees Celsius, with bars indicating standard error from triplicate samples. Replicate samples from WT-1 and WT-2 are combined and reported jointly as WT. Samples are shown in order from left to right: WT, Q22T, F155G, W123R, V401A, Y169M, W438D.

### Comparison between thermal stability results and initial Foldit predictions

In Foldit,^7^ the predicted changes in stability of an enzyme is evaluated by the calculated energy, measured with a negative number if a change is predicted to be stabilizing; positive changes occur when predicted to be destabilized. When comparing the Foldit free energy predictions with the experimental changes in mean thermal stability, there appears to be a modest positive correlation (Pearson’s Correlation = 0.77). However, this data will need to be evaluated in the context of additional mutations and larger datasets with forward engineering efforts to determine if this trend is more than a spurious association.

**Figure 6.**
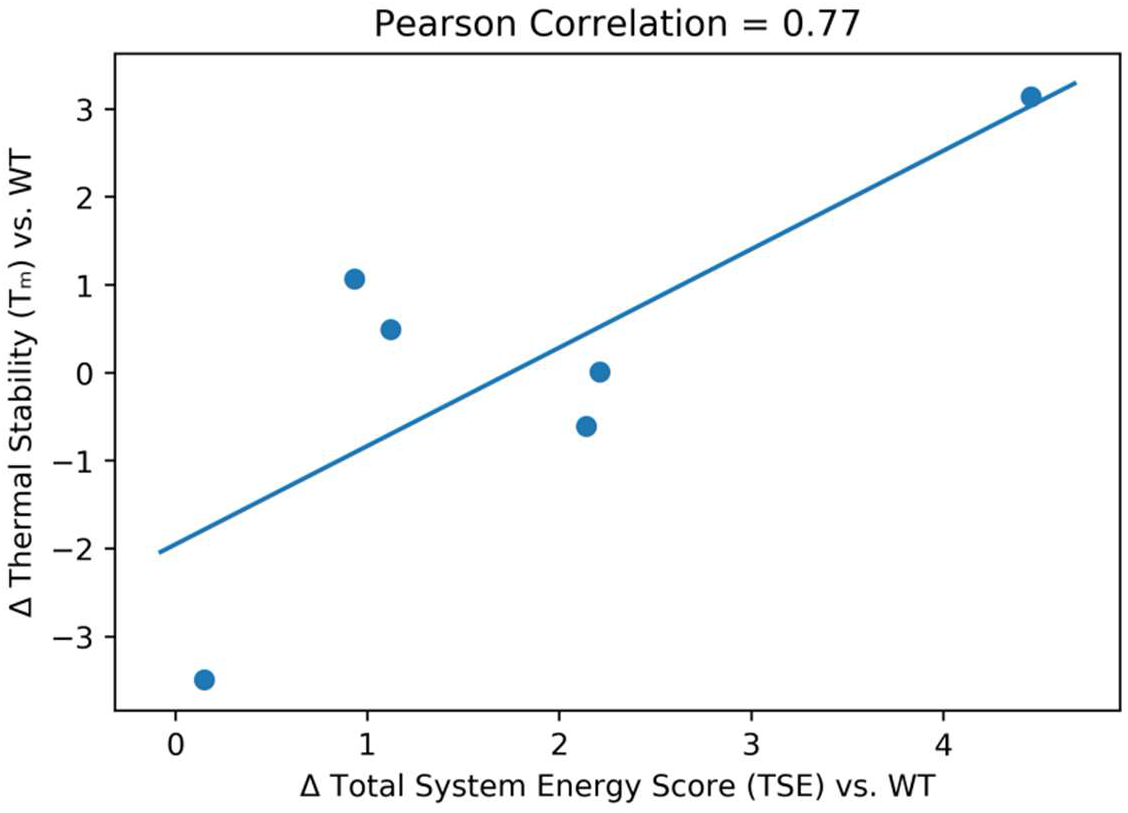
Foldit score changes vs. Experimental change in thermal stability. Changes in T_M_ values compared to the wildtype are plotted in units of degrees Celsius in the vertical axis while changes in TSE compared to the wildtype are in the horizontal axis.

## Discussion

In this study, six variants of β-Glucosidase B in select regions were designed and experimentally analyzed for catalytic proficiency and thermal stability. Two wildtype BglB samples were produced and tested alongside the variants. All six variants designed in this study were found able to express and be isolated as a soluble protein and therefore were evaluated for their catalytic and thermal stability parameters.

For the variant Q22T, we observed a significant plummet in the *k*_cat_ value (p < 0.01) while the K_M_ remained consistent with the native BglB. Before the mutation, Foldit suggested a hydrogen bond is formed between the glucose and the original glutamine, located 3Å apart. As the glutamine is mutated to threonine, the distance increases, and Foldit predicted the hydrogen bond would be disrupted. The loss in a key interaction that likely stabilizes substrate binding and transition state formation may explain the decrease in catalytic activity.

Variant Y169M exhibited a threefold increase of K_M_, indicating that the mutation had changed enzyme-substrate interactions. It appeared that the original tyrosine may play a role in hydrogen bonding interactions with the substrate. The change from tyrosine to methionine may cause difficulties for the substrate to reach the active site due to losing potential factors such as pi-pi stacking. The thermal stability of Y169M remained unchanged from that of the WT and we hypothesize that this may be due to the lack of hydrogen bonding in native tyrosine.

Variant W438D saw reductions in catalysis while maintaining a similar thermal stability as the wild type. Compared to the wild type, its *k*_cat_/K_M_ value exhibited an eight-fold reduction. This data contradicts the notion that mutations on the periphery of the enzyme will have little impact on catalysis, since its location is far from the active site. The cause for this decrease in kinetic parameters is unknown. Experimental errors such as possible contamination may have occurred, and more replications should be performed on all variants in this study by future experiments to reproduce the results.

Thermal stability results indicated an overall decrease in the thermal stability between the six variants, with only W123R exhibiting increased T_M_. This is in accordance with the initial hypothesis. Despite all mutants exhibiting a decrease in kinetic activity by more than two-fold, kinetic parameters fluctuated in association with variant-specific physico-chemical changes, such as hydrogen bonding and electrostatic attraction.

Variant W123R resulted in an increase in thermal stability, raising T_M_ by more than 3 °C. Surprisingly, this was not predicted by Foldit: TSE score of W123R is 4.45 units higher than the wild type, which points to decreased thermal stability. Physio-chemical analysis indicate that the mutation to a positively charged arginine can result in new, strong electrostatic interactions with the neighboring glutamate 164, which increases thermal stability.

Variant F155G exhibited relatively similar thermal stability to the wild type but with a halved kinetic activity. The mutation from a large, nonpolar Phenylalanine to a smaller Glycine offers more flexibility and unpredictability in enzyme-substrate interactions, which may have resulted in the decrease in kinetic activity.

For Variant V401A, Foldit predicted a similar TSE score; however, there was a significant decrease in T_M_ (p < 0.01). Prior to the mutation, the side chain of valine 398 is hydrophobic. As the valine is mutated to a smaller alanine, pockets that allows water to enter the active site may be created, which can disrupt its overall thermal stability.

The linear regression analysis of total system energy and thermal stability contradicts with respect to the hypothesized changes in the variants’ thermal stability, since increases in total system energy score is associated with decreased molecular stability. This finding points to limitations of modeling tools’ power for predicting enzymatic functional parameters. While this result aligns with some of the more extensive work of Carlin et al 2016 and Carlin et al 2017, given the sample size, the observable trend may be a result of spurious correlation.

Collectively, the results described here highlight the complexity of protein sequence structure-function space and the need for the continued expansion of data sets. These results have been integrated into a larger data set that maintains the high resolution of quantitative findings presented here, which is being developed to train more advanced protein prediction algorithms.

---

**SI Table 1.** Total System Energy Score, Expression, *k*_cat_, K_M_, *k*_cat_/K_M_, T_M_ of six variants and two wildtypes

| | TSE | Expression<br>(mg/ml) | $k_{\text{cat}}$ ($\text{min}^{-1}$) | $K_M$ (mM) | $k_{\text{cat}} / K_M$ ($\text{min}^{-1}\text{mM}^{-1}$) | $T_M$ ($^{\circ}\text{C}$) |
| --- | --- | --- | --- | --- | --- | --- |
| WT-1 | -544.36 | 2.29 | $37.19 \pm 3.36$ | $5.4 \pm 0.88$ | $6.89 \pm 1.28$ | $44.9 \pm 0.12$ |
| Q22T | -543.24 | 0.29 | $0.26 \pm 0.06$ | $99.02 \pm 247.88$ | $0.0026 \pm 0.01$ | $44.68 \pm 0.60$ |
| W123R | -539.91 | 1.49 | $0.015 \pm 0.08$ | $49.63 \pm 324.02$ | $0.00020 \pm 0.01$ | $47.33 \pm 0.76$ |
| F155G | -543.43 | 0.56 | 18.56 | 6.10 | 3.04 | $45.26 \pm 0.15$ |
| WT-2 | -544.36 | 2.29 | $47.21 \pm 3.36$ | $5.98 \pm 0.88$ | $7.89 \pm 1.29$ | $44.19 \pm 0.07$ |
| W438D | -542.16 | 0.84 | $10.59 \pm 1.65$ | $10.28 \pm 2.14$ | $1.03 \pm 0.27$ | $44.2 \pm 0.02$ |
| Y169M | -542.22 | 2.53 | $50.57 \pm 13.63$ | $22.2 \pm 5.53$ | $2.28 \pm 0.84$ | $43.58 \pm 0.26$ |
| V401A | -544.22 | 0.31 | $20.46 \pm 1.29$ | $8.27 \pm 1.24$ | $2.47 \pm 0.40$ | $40.7 \pm 0.13$ |

The estimated errors are calculated with standard deviation of three independent measurements.

